# Cerebral blood flow predicts multiple demand network activity and fluid intelligence across the lifespan

**DOI:** 10.1101/2021.11.10.468042

**Authors:** Shuyi Wu, Lorraine K. Tyler, Richard N.A. Henson, James B. Rowe, Cam-Can, Kamen A. Tsvetanov

## Abstract

The preservation of cognitive function into old age is a public health priority. Cerebral hypoperfusion is a hallmark of dementia but its impact on maintaining cognitive ability across the lifespan is less clear. We investigated the relationship between baseline cerebral blood flow (CBF) and blood oxygenation level-dependent (BOLD) response during a fluid reasoning task in a population-based adult lifespan cohort (N=227, age 18-88 years). As age differences in baseline CBF could lead to non-neuronal contributions to the BOLD signal, we introduced commonality analysis to neuroimaging, in order to dissociate performance-related CBF effects from the physiological confounding effects of CBF on the BOLD response. Accounting for CBF, we confirmed that performance- and age-related differences in BOLD responses in the multiple-demand network (MDN) implicated in fluid reasoning. Differences in baseline CBF across the lifespan explained not only performance-related BOLD responses, but also performance-independent BOLD responses. Our results suggest that baseline CBF is important for maintaining cognitive function, while its non-neuronal contributions to BOLD signals reflect an age-related confound. Maintaining perfusion into old age may serve to support brain function with behavioural advantage, regulating brain health.

## 1. Introduction

The world’s population is ageing, with every sixth person expected to be over 65 by 2050 (United Nations, 2020). Cognitive decline has emerged as a major health threat in old age, including but not limited to dementia (Piguet et al., 2009; Yarchoan et al., 2012). To combat this threat, there is increasing demand to identify factors that facilitate the maintenance of cognitive function across the lifespan. Ageing causes changes to our brains in vascular, structural and functional domains (Kennedy and Raz, 2015; Cabeza et al., 2018). However, these effects are normally reported separately, and only through their integration one can better understand how these domains influence cognitive decline in old age (Tsvetanov et al., 2021).

Cerebral blood flow (CBF) changes early in experimental models of dementia, leading to neuronal dysfunction, and loss independently of amyloid-β-dependent contributions (Iadecola, 2004; Zlokovic, 2011; Kisler et al., 2017; Sweeney et al., 2018, 2019). In healthy ageing, previous reports have linked the effects of age on baseline CBF to behavioural performance measured outside of the scanner (Bangen et al., 2014; Hays et al., 2017; Leeuwis et al., 2018). However, brain perfusion measurements are highly dependent on other physiological factors such as autoregulation modulators (Lemkuil et al., 2013), medication, time of day, levels of wakefulness (Patricia et al., 2014), physical exercise, caffeine or smoking before the scan (Domino et al., 2004; Addicott et al., 2009; Merola et al., 2017). Therefore, differences in CBF signal may reflect an age-related bias in such factors, rather than a true baseline difference in CBF (Grade et al., 2015). Moreover, it remains unclear whether the observed CBF dysregulation in ageing reflects a link between somatic differences in vascular health and global cognition, or whether CBF modifies regional brain activations underlying specific cognitive processes. To understand the role of baseline CBF in cognitive ageing, one must also test whether baseline CBF is associated with performance-related brain activity during cognitive tasks.

The field of neurocognitive ageing research has often used functional magnetic resonance imaging (fMRI) to study age differences in brain activity during cognitive tasks. FMRI data are usually interpreted in terms of neuronal activity, but the blood oxygenation level-dependent (BOLD) signal measured by fMRI also reflect vascular differences and neurovascular coupling (Mishra et al., 2021), which changes with age (Tsvetanov et al., 2021). Failure to account for vascular health alterations leads to misinterpretation of fMRI BOLD signals (Hutchison et al., 2013; Liu et al., 2013; Tsvetanov et al., 2015) and their cognitive relevance (Geerligs and Tsvetanov, 2016; Tsvetanov et al., 2016; Geerligs et al., 2017). Several approaches exist to separate vascular from neural contributions to the BOLD signals, including the use of baseline CBF to normalise for age differences in cerebrovascular function (Tsvetanov et al., 2021). Normalisation with baseline CBF would improve detection of “true” neuronal changes i.e., over and above age-related differences in non-neuronal physiology. This would control for behaviourally irrelevant *confounding effects,* and *performance-related effects* where cerebral hypoperfusion reflects neuronal function and loss. Therefore, it would be better to integrate, not simply control for, baseline CBF differences in task-based BOLD studies to dissociate confounding from performance-related effects of CBF on age-related differences in the BOLD fMRI responses.

To distinguish confounding from performance effects of CBF is important to understand the neuronal substrates of multiple cognitive demands with ageing (Kaufman and Horn, 1996; Salthouse, 2012; Kievit et al., 2014). Demanding, complex or executive functions depend on a distributed network of brain regions known as the multiple-demand network (MDN), which is readily activated during tasks used to assess fluid intelligence (Crittenden et al., 2016; Tschentscher et al., 2017; Woolgar et al., 2018). The MDN parses complex tasks into subcomponents or sub-goals (Duncan, 2013; Camilleri et al., 2018). There is substantial spatial overlap between MDN and the brain regions with impaired baseline CBF in ageing (Tsvetanov et al., 2020b, 2021). Therefore, some of the age differences in MDN and cognition (Tsvetanov et al., 2016; Samu et al., 2017) may reflect confounding and/or performance-related effects of CBF dysregulation.

To characterise neurocognitive ageing, we propose the use of commonality analysis to dissociate *confounding* from *performance-related* effects of CBF on age-related differences in brain functional measures. Commonality analysis, unlike the normalisation approach, allows for adjustment of multiple variables simultaneously by identifying the variance in a dependent variable associated with each predictor uniquely, as well as the variance in common to two or more predictors (Nimon et al., 2008; Kraha et al., 2012). Here, we identify unique and common effects of age, performance, and baseline CBF on fMRI BOLD responses during a fluid reasoning task in a population-based adult lifespan cohort (age 18-88, N = 227, www.camcan.org). Reasoning was measured by the common Cattell task of fluid intelligence, which requires solving a number of problems, and is known to decline dramatically with age (Kievit et al., 2014).

We predicted that the integration of baseline CBF with task-based fMRI BOLD would improve detection of confounding and performance-related effects of CBF associated with reasoning. Performance-related effects of CBF would be indicated by variance in the BOLD response that is common to age, task performance and CBF, whereas confounding effects of CBF would be indicated by variance that is common to age and CBF, but not shared with performance.

## 2. Methods

### 2.1. Participants

Figure 1 illustrates the study design, data processing and analysis pipeline. The data were acquired from Phase 3 of the Cambridge Centre for Aging and Neuroscience (Cam-CAN), a large population-based study of the healthy adult life span (Shafto et al., 2014; Taylor et al., 2015). The ethical approval for the study was approved by the Cambridge 2 Research Ethics Committee and written informed consent was provided by all participants. Exclusion criteria included poor hearing (a sensitive threshold of 35 dB at 1000 Hz in both ears) and poor vision (below 20/50 on the Snellen test; Snellen, 1862), low Mini-Mental Status Examination (Folstein et al., 1975), self-reported substance abuse as assessed by the Drug Abuse Screening Test (Skinner, 1982), significant psychiatric disorders (e.g., schizophrenia, bipolar disorder, personality disorder), or neurological diseases (e.g. a history of stroke, epilepsy, traumatic brain injury). Demographic characteristics of the sample are described in Table 1.

**Figure 1.**
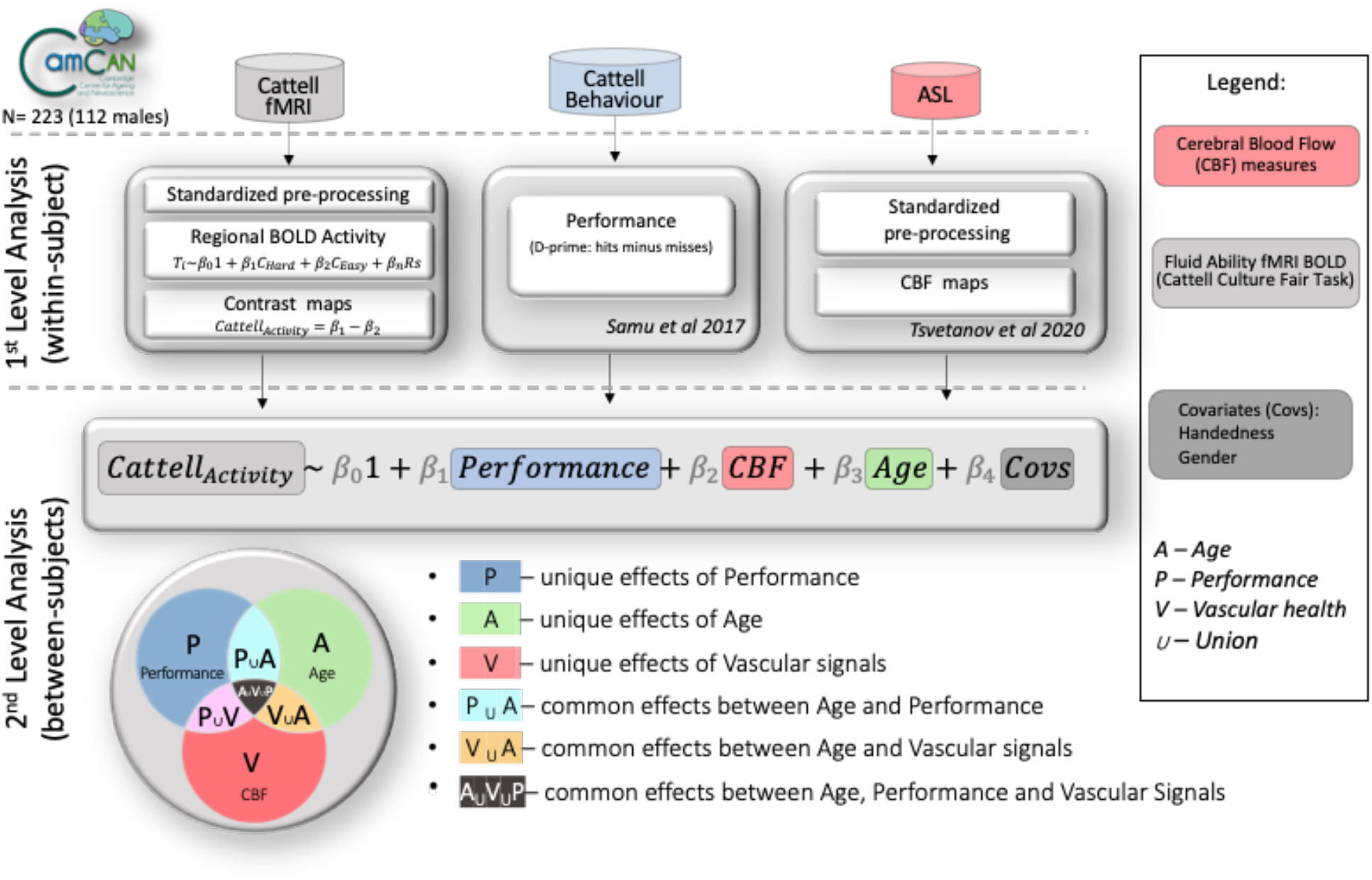
Summary of the analytical pipeline.

**Table 1.** Participants’ demographic information

| N=223 | Decile |  |  |  |  |  |  |
| --- | --- | --- | --- | --- | --- | --- | --- |
|  | 1 | 2 | 3 | 4 | 5 | 6 | 7 |
| Age range [years] | 19-27 | 28-37 | 38-47 | 48-57 | 58-67 | 68-77 | 78-87 |
| Numbers | 21 | 39 | 37 | 35 | 35 | 30 | 26 |
| Gender |  |  |  |  |  |  |  |
| Male | 9 | 19 | 18 | 17 | 18 | 16 | 14 |
| Female | 12 | 20 | 19 | 18 | 17 | 14 | 12 |
| Handedness<sup>a</sup> |  |  |  |  |  |  |  |
| Mean/SD | 79/44 | 89/25 | 81/27 | 94/11 | 77/50 | 95/10 | 87/33 |
| Range[min/max] | -100/100 | -56/100 | -56/100 | 58/100 | -78/100 | 53/100 | -56/100 |
| Education<sup>b</sup> |  |  |  |  |  |  |  |
| None | 0 | 0 | 0 | 0 | 0 | 5 | 1 |
| GCSE | 2 | 2 | 6 | 3 | 3 | 3 | 3 |
| A-level | 4 | 1 | 3 | 11 | 9 | 8 | 9 |
| University | 15 | 36 | 28 | 21 | 23 | 14 | 13 |

<sup>a</sup> Higher scores indicate greater right-hand preference, as assessed by Edinburgh Handedness Inventory (Oldfield, 1971).
<sup>b</sup> Categorized according to the British education system: "None" = no education over the age of 16 yrs; "GCSE" = General Certificate of Secondary Education; "A Levels" = General Certificate of Education Advanced Level; "University" = undergraduate or graduate degree.

### 2.2. Stimuli, task and procedure

Participants undertook a Fluid Intelligence task which draws on critical cognitive process of fluid reasoning, which underlies many complex cognitive operations (Duncan, 2013), and which declines with age (Horn and Cattell, 1967; Kaufman and Horn, 1996; Salthouse et al., 2003; Duncan, 2010; Salthouse, 2012; Kievit et al., 2014). We used a simplified version of the Cattell Culture Fair test (Cattell, 1971), modified to be used in the scanner (Woolgar et al., 2013; Samu et al., 2017). On each trial, participants were presented with a display of four patterns and had to select the “odd one out”. The task employed a block design, with 30-seconds blocks of trials alternating between two conditions with different difficulty level (“easy” and “hard” puzzles). There was a total of four blocks per condition. Because there was a fixed time to perform as many trials as possible, behavioural performance was measured by subtracting the number of incorrect trials from the number of correct trials (averaged over hard and easy bocks, following Samu et al., (2017), i.e. to ensure that someone responding quickly but randomly did not score highly. The suitability of this performance score was confirmed by its strong correlation (Pearson’s r[95% CI]: r(223) = 0.70 [0.63, 0.76], *P* < 0.001) with scores obtained from the full version of the Cattell test, administered outside the scanner at stage 2 of Cam-CAN (Shafto et al., 2014). We also excluded n = 28 participants who had disproportionately poor performance with 10 or more incorrect trials (17 females, with age range 31-88); leaving N = 223 remaining (111 females, age range 19 - 87 years).

### 2.3. MRI Acquisition and Preprocessing

Imaging data were acquired using a 3T Siemens TIM Trio System with a 32-channel head-coil at the MRC Cognition and Brain sciences Unit (CBU; www.mrc-cbu.cam.ac.uk). Of the initial cohort, 256 participants had valid T1, T2, arterial spinning labelling (ASL) data, and task-induced BOLD data from a fluid intelligence task.

A 3D-structural MRI was acquired on each participant using T1-weighted sequence (Generalized Auto-calibrating Partially Parallel Acquisition (GRAPPA) with the following parameters: repetition time (TR) = 2,250 ms; echo time (TE) = 2.99 ms; inversion time (TI) = 900 ms; flip angle α= 9°; field of view (FOV) = 256 × 240 × 192 mm^3^; resolution = 1 mm isotropic; accelerated factor = 2; acquisition time, 4 min and 32 s.

We used Release003 of the CamCAN Automatic Analysis pipelines for Phase III data (Taylor et al.,(2015), which called functions from SPM12 (Wellcome Department of Imaging Neuroscience, London, UK). The T1 image from Phase II was rigid-body coregistered to the MNI template, and the T2 image from Phase II was then rigid-body coregistered to the T1 image. The coregistered T1 and T2 images were used in a multimodal segmentation to extract probabilistic maps of six tissue classes: gray matter (GM), white matter (WM), CSF, bone, soft tissue, and residual noise. The native space GM and WM images were submitted to diffeomorphic registration to create group template images. Each template was normalized to the MNI template using a 12-parameter affine transformation.

### 2.4. EPI image acquisition and processing

For the Cattell-based fMRI in Phase III of CamCAN, Gradient-Echo Echo-Planar Imaging (EPI) of 150 volumes captured 32 axial slices (sequential descending order) of thickness of 3.7 mm with a slice gap of 20% for whole-brain coverage with the following parameters: TR = 1970 ms; TE = 30 ms; flip angle α = 78°; FOV = 192 × 192 mm^2^; resolution = 3 × 3 × 4.44 mm^3^, with a total duration of 5 min.

EPI data preprocessing included the following steps: (1) spatial realignment to adjust for linear head motion, (2) temporal realignment of slices to the middle slice, (3) coregistration to the T1 anatomical image from Phase II above, (4) application of the normalization parameters from the T1 stream above to warp the functional images into MNI space, and (5) smoothing by an 8mm Gaussian kernel.

For the participant-level modelling, every voxel’s time-course was regressed in a multiple linear regression on the task’s design matrix which consisted of time-courses for hard and easy conditions convolved with a canonical haemodynamic response function (HRF). Regressors of no interest included WM, CSF, 6 standard realignment parameters (accounting for in-scanner head motions), and harmonic regressors that capture low-frequency changes (1/128 Hz) in the signal typically associated with scanner drift and physiological noise. WM and CSF signals were estimated for each volume from the mean value of WM and CSF masks derived by thresholding SPM’s tissue probability maps at 0.75. The contrast of parameter estimates for hard minus easy conditions for each voxel and participant was then calculated, termed here *Cattell activation*.

### 2.5. Arterial spinning labelling (ASL) image acquisition and processing

Perfusion-weighted images of cerebral blood flow used pulsed arterial spin labelling (PASL, PICORE-Q2T-PASL with background suppression). The sequence is used with the following parameters: repetition time (TR) = 2500 ms, echo time (TE) = 13 ms, field of view (FOV) = 256 × 256 × 100 mm^3^, 10 slices, 8 mm slice thickness, flip angle = 90°, inversion time 1 (TI1) = 700 ms, TI2 = 1800 ms, Saturation stop time = 1600 ms, tag width = 100 mm and gap = 20.9 mm, 90 repetitions giving 45 control-tag pairs, voxel-size = 4 mm × 4 mm × 8 mm, 25% interslice gap, acquisition time of 3 minutes and 52 seconds. In addition, a single-shot EPI (M0) equilibrium magnetization scan was acquired. Pulsed arterial spin labelling time series were converted to maps of CBF using Explore ASL toolbox (https://github.com/ExploreASL/ExploreASL; Mutsaerts et al., 2018). Following rigid-body alignment, the images were coregistered with the T1 from Phase II above, normalised with normalization parameters from the T1 stream above to warp ASL images into MNI space and smoothed with a 12 mm FWHM Gaussian kernel (for more details, Tsvetanov et al., 2020b).

### 2.6. Analytical approach

To model random effects across participants, we performed voxel-wise analysis using multiple linear regression (MLR) with age as the main independent variable of interest, and sex and handedness as covariates of no interest. This MLR was applied to maps of both Cattell activation (BOLD) and baseline CBF.

To evaluate the confounding and performance-related effects of resting CBF on BOLD activation, we conducted commonality analysis (Nimon et al., 2008; Kraha et al., 2012). Commonality analysis partitions the variance explained by all predictors in MLR into variance unique to each predictor and variance shared between each combination of predictors. Therefore, unique effects indicate the (orthogonal) variance explained by one predictor over and above that explained by other predictors in the model, while common effects indicate the variance shared between correlated predictors. Notably, the sum of variances, also known as commonality coefficients, equals the total R^2^ for the regression model.

We adapted a commonality analysis algorithm (Nimon et al., 2008) for neuroimaging analysis to facilitate voxel-wise nonparametric testing in Matlab (Mathworks, https://uk.mathworks.com/). The commonality analysis was applied the Cattell activation in each voxel separately (see Figure 1). The independent variables in the model were baseline CBF for the corresponding voxel, age and task performance. Covariates of no interest included sex and handedness. The model can therefore identify unique variance explained by each of the predictors (U_CBF_, U_Age_ and U_P_ for CBF, Age and Performance, respectively). Common effects of interest were the *confounding effects*, defined by the shared variance between CBF and age (C_CBF,Age_), and *performance-related effects*, defined by the common variance between CBF, Age and Performance (C_CBF,Age,P_). Significant clusters related to effects of interest were identified with nonparametric testing using 1000 permutations and threshold-free cluster enhancement, corrected to p<.05 (Smith and Nichols, 2009). This Matlab version of commonality analysis for neuroimaging with TFCE implementation is available at https://github.com/kamentsvetanov/CommonalityAnalysis/.

### 2.7. Data and code availability

The dataset analysed in this study is part of the Cambridge Centre for Ageing and Neuroscience (Cam-CAN) research project (www.cam-can.com). Raw and minimally pre-processed MRI (i.e. from automatic analysis; Taylor et al., 2015) and behavioural data are available by submitting a data request to Cam-CAN (https://camcan-archive.mrc-cbu.cam.ac.uk/dataaccess/).

Task-based fMRI data was post-processed using SPM12 (http://www.fil.ion.ucl.ac.uk/spm; Friston et al., 2007). Arterial spin labelling data were post-processed using ExploreASL toolbox (https://github.com/ExploreASL/ExploreASL; Mutsaerts et al., 2018). As part of this study, MATLAB-based commonality analysis for neuroimaging with TFCE implementation was developed and made available at https://github.com/kamentsvetanov/CommonalityAnalysis/. Visualisation of all neuroimaging results was generated using MRIcroGL (https://github.com/rordenlab/MRIcroGL; Rorden and Brett, 2000). The corresponding author (K.A.T.) can provide custom-written analyses code on request.

## 3. Results

### 3.1. Main effect and effect of age on BOLD in Cattell task

Group-level analysis confirmed activations for the hard vs easy condition in the lateral prefrontal cortex, the anterior insula, the dorsal anterior cingulate cortex, the frontal eye field, the pre-supplementary motor area and areas along the intraparietal sulcus and lateral temporal lobe, recapitulating the multiple demand network (MND, Duncan, 2013; Camilleri et al., 2018), and the lateral occipital cortex and the calcarine cortex (Figure *2*a). Additionally, we observed deactivations in the ventral medial prefrontal cortex (vmPFC), posterior cingulate cortex (PCC) and inferior parietal lobe (IPL), recapitulating the default network (Buckner et al., 2008; Raichle, 2015; Buckner and DiNicola, 2019). With respect to ageing, there were weaker activations in regions of the MDN, and weaker deactivations in regions of the DMN, associated with increasing age, see Figure *2*b, consistent with previous studies (Samu et al., 2017).

**Figure 2.**
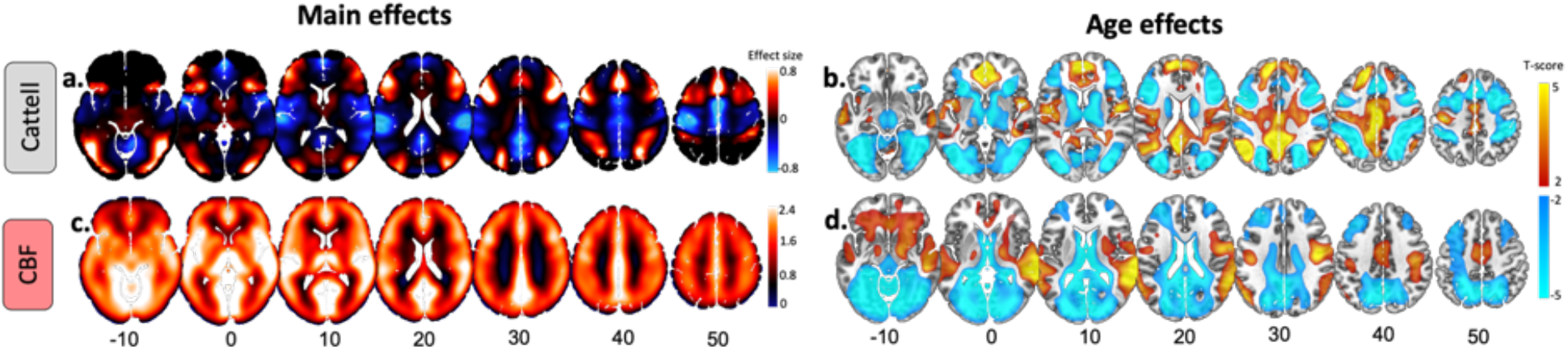
Main and age effects on task-based activity and cerebral blood flow (CBF) maps. (a) Main effects of BOLD activity in response to Hard vs Easy blocks with over- and under-activations shown in warm and cold colours, respectively. (b) Age-related decreases (cold colours) and increases (warm colours) in Cattell task. (c) Main effect of baseline CBF across all participants. (d) Age-related decreases (cold colours) and increases (warm colours) in baseline CBF. Slices are numbered by z level in Montreal Neurological Institute (MNI) space.

### 3.2. Main effect and effect of age on baseline CBF

Group-level results revealed a pattern of relatively high cerebral blood flow in cortical and subcortical brain areas associated with high perfusion and high metabolism (Henriksen et al., 2018; Figure 2c), such as caudal middle-frontal, posterior cingulate, pericalcarine, superior temporal and thalamic regions. Moderate to low CBF values in the superior-parietal and inferior-frontal areas of the cortex (Figure *2*c) may reflect the axial positioning of the partial brain coverage sequence used in the study.

We observed age-related declines in CBF in the bilateral dorsolateral prefrontal cortex, lateral parietal cortex, anterior and posterior cingulate, pericalcarine, and cerebellum (Figure 2c) in agreement with previous reports (Chen et al., 2011; Zhang et al., 2018). Also, we observed age-related CBF increase in regions susceptible to individual and group differences in arterial transit time that can bias accuracy of CBF estimation, including middle temporal gyrus and middle cingulate cortex (Mutsaerts et al., 2017).

### 3.3. Commonality analysis of BOLD Cattell activation

#### 3.3.1. Unique effects

Unique effects of individual differences in performance levels on Cattell activation (BOLD) were found in regions similar to those activated by the main effect of the Cattell task (e.g. MDN), with the exception of the lateral occipital cortex and inclusion of inferior temporal gyrus, primary visual cortex, caudate and thalamus, (cf. Figure 2a and Figure 3 top panel). Unlike the case for main effects, task-negative regions (e.g., DMN) showed small to no significant associations with performance.

**Figure 3.**
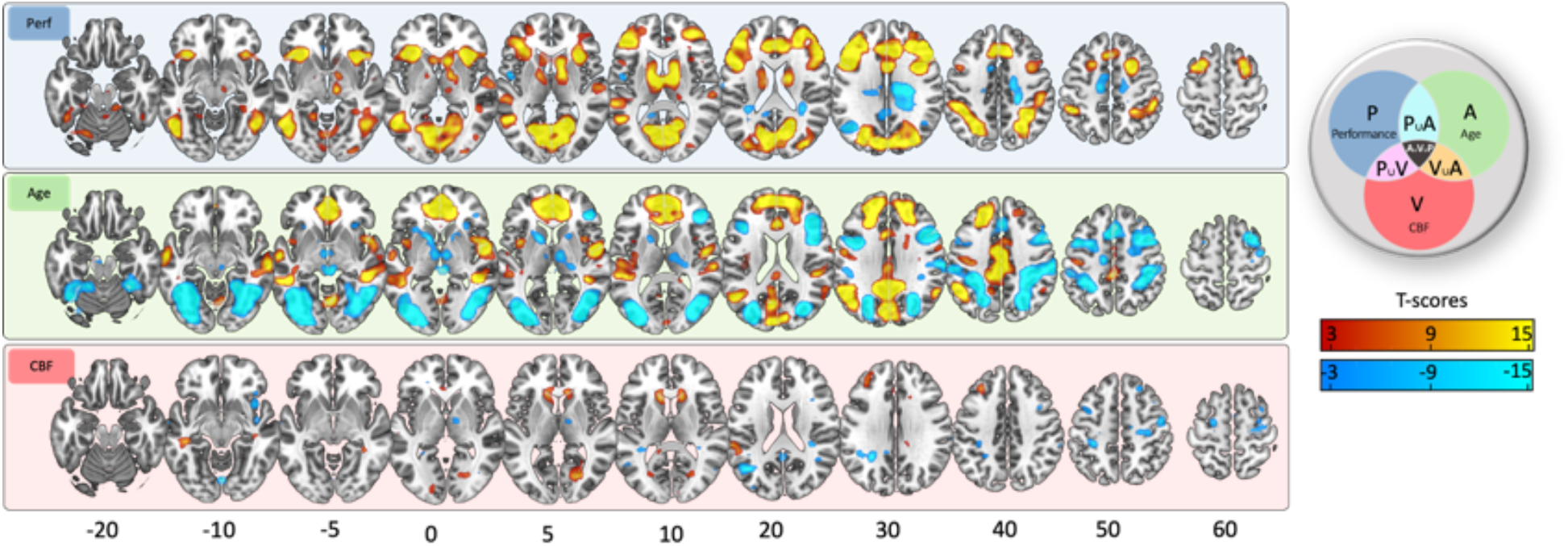
Unique effects in commonality analysis. (top panel) Age-related decreases (cold colours) and increases (warm colours). (middle panel) Performance-related decreases (cold colours) and increases (warm colours). (bottom panel) CBF-related decreases (cold colours) and increases (warm colours) in Cattell task. Slices are numbered by z level in Montreal Neurological Institute (MNI) space.

Unique effects of age were similar but weaker to the effect of age in the model without other predictors (cf. Figure *2*b and Figure 3 middle panel). Unique positive associations between CBF and Cattell activation was observed in middle frontal gyrus and cuneus regions. Negative associations were observed in insular regions, posterior cingulate cortex, bilateral angular gyrus, precentral gyrus and superior frontal gyrus.

Unique effects of CBF were weak, but significant, showing positive associations with activation in the middle frontal gyrus, the putamen and the cuneus. Additionally, CBF was associated negatively with activation in task negative regions, namely the angular gyrus and precentral gyrus.

#### 3.3.2. Common effects

There were many common effects between age and performance, with a positive commonality coefficient (C_Age,P_, Figure 4, cyan colour), i.e. a portion of the age effects on Cattell activation was related to performance effects on Cattell activation. These effects were observed in task positive (e.g. MDN) and task negative regions (e.g. DMN), in addition to thalamus, caudate, primary motor cortex.

**Figure 4.**
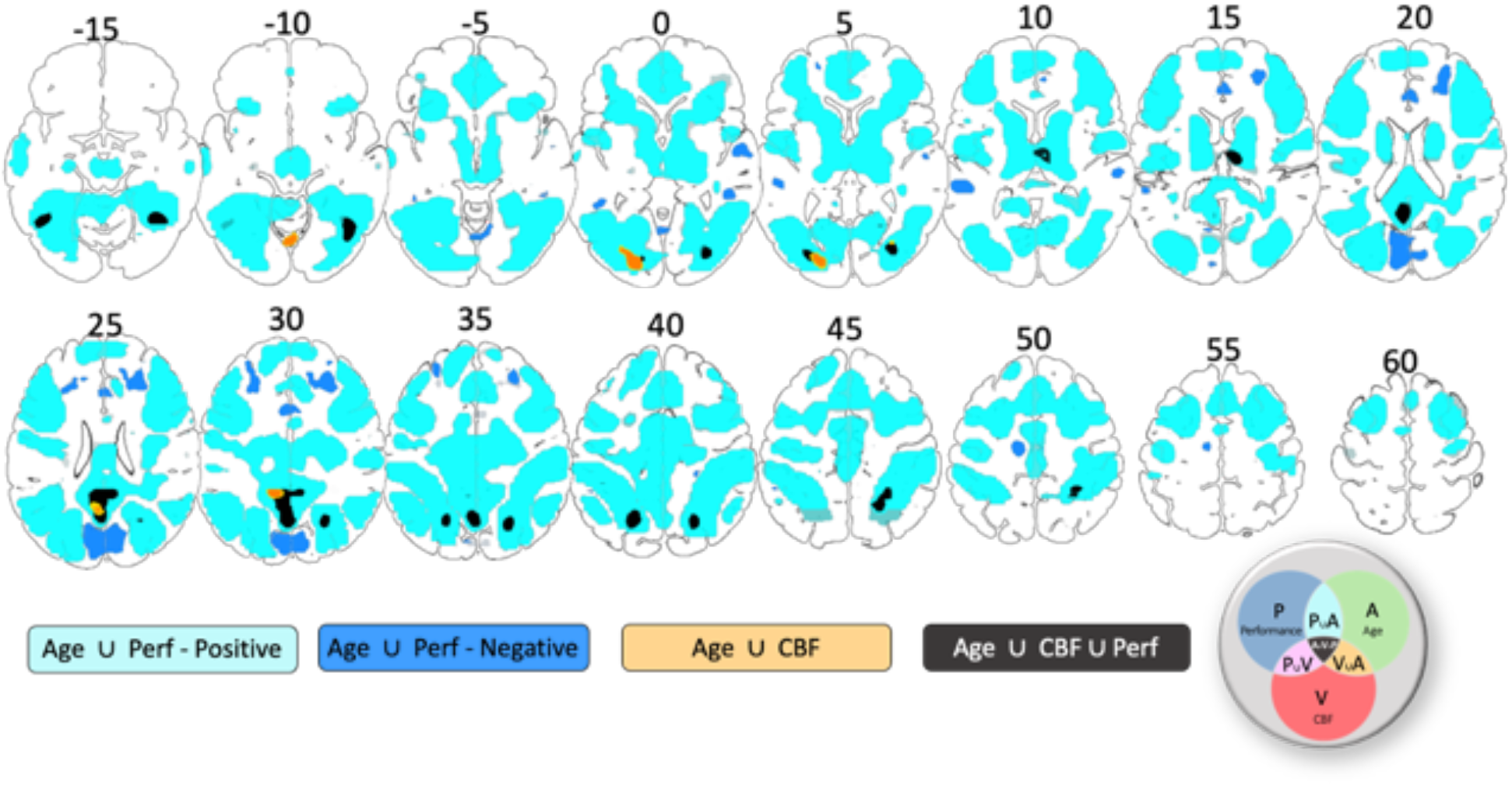
Common Effects in commonality analysis. Positive and negative common effects between age and performance are shown in cyan and dark blue colours, respectively. Common effects between age and baseline CBF are shown in orange colour. Common effects between age, performance and CBF are shown in black colour. P – performance, A – age, V – vascular, i.e. CBF. Slices are numbered by z level in Montreal Neurological Institute (MNI) space.

Negative commonality coefficients between performance and age were observed in the cuneus, bilateral middle frontal gyrus, anterior and middle cingulate gyrus, and bilateral superior temporal gyrus (dark blue colour in Figure 4). Negative values of commonality coefficients indicate a suppressor relationship between predictors (Zientek and Thompson, 2006), i.e. the effects of age and/or performance are stronger with their joint consideration in the model.

Confounding effects of baseline CBF on Cattell activation were characterised by the common effect between Age and CBF (C_CBF,Age_). Significant confounding effects were localised within posterior cingulate cortex, fusiform gyrus and inferior occipital gyrus (orange colour in Figure 4).

Performance-related effects of baseline CBF on Cattell activation were characterised by the common effect between Age, CBF and Performance (C_CBF,Age,P_, black colour in Figure 4). Regions included intraparietal sulcus, posterior cingulate cortex, precuneus, thalamus and fusiform gyrus. Furthermore, consistent with previous findings (Tsvetanov et al., 2018), behaviourally-relevant effects were seen in inferior temporal and adjacent occipital regions, presumably due to attentional enhancement of visual representations in the more difficult conditions (Fedorenko et al., 2013).

## 4. Discussion

The study confirmed the prediction that regional cerebral blood flow (CBF) can explain both performance-related and age-dependent components of the fMRI BOLD signal in parts of the multiple-demand network (MDN) associated with more complex reasoning during a common test of fluid intelligence (Cattell task). The age-dependent differences in baseline CBF also explained variance in fMRI BOLD signal in some regions that was not related to task-performance. We propose that modelling the effects of age on baseline CBF, and in general cerebrovascular and neurovascular health (Tsvetanov et al., 2021), improves the interpretation of fMRI studies, with implications for understanding brain health with ageing and disease, and that maintaining brain perfusion as we get older may have a protective effect on brain function and cognition.

### 4.1. Age differences in baseline cerebral blood flow are related to behaviour-relevant Cattell BOLD activity

Age-related decreases in baseline cerebral blood flow (CBF), assessed with a non-invasive MR-perfusion technique, related to behaviourally relevant BOLD activity evoked by demanding problem-solving. Our findings are consistent with previous studies relating baseline CBF to performance on tasks carried outside the scanner (Bangen et al., 2014; Hays et al., 2017). We extend these lines of work by showing that baseline CBF is linked to BOLD activity, with behavioural correlation across individuals. Age-related decrease in CBF and decline in performance related to a lower range of activation in task-positive regions and less deactivation of task-negative regions. Of all task-positive regions, the bilateral intra-parietal sulcus, the thalamus, and the fusiform gyrus showed significant common effects between age, CBF and performance. The intraparietal sulcus and the thalamus also showed a unique association between performance and BOLD activity, suggesting a neural origin of the effects in these regions. The processes contributing to coupling between baseline CBF and neural activity are multifaceted, probably comprising neurogenic vasodilation, cardiac output and arterial remodelling (Gaballa et al., 1998; Ohanian et al., 2014; Li et al., 2015), all of which change with age and regulate baseline and stimulus-evoked CBF (Willie et al., 2014). Establishing the relative contribution and importance of these processes warrants future research.

Of all task-negative regions, only the posterior cingulate cortex showed common effects between age, CBF and performance in predicting BOLD activation in the Cattell task. In this region, age-related reduction in CBF and performance correlated with less deactivation in the posterior cingulate cortex. The posterior cingulate cortex did not show unique effects between performance and BOLD activity, suggesting a mechanism different from the one observed in task-positive regions, likely reflecting a non-neuronal origin of the effects (see also “Unique effects of performance, age and CBF in Cattell task”). While the deactivation of the default network in young adults is thought to reflect suppression of neuronal activity (Fox et al., 2018), in the present study, some of the poor performing older adults showed an over-activation, not less deactivation. This again suggests a different involvement of the posterior cingulate cortex in older adults compared to young adults, for instance, signals of non-neuronal origin caused by physiological artifacts (Birn et al., 2006; Tsvetanov et al., 2021) or ‘vascular steal’ (Shmuel et al., 2002). Taken together, these findings may reflect compromised vasodilatory reserve, resulting in an inefficient redirection of resources from task-positive regions to task-negative regions in the attempt to meet higher energy demands in task-positive regions, perhaps reflecting blood flow-dependent glycolysis and oxidative metabolism. The breath of these associations is consistent with theories of vasoactive and cardiovascular regulation of cerebral blood flow (Sobczyk et al., 2014; Digernes et al., 2017).

### 4.2. Vascular Confounding effects of CBF on task-related activity

Only a portion of the age differences in performance-independent BOLD activation were associated CBF decreases. Furthermore, the effects were observed in non-classical demand network task-positive and task-negative regions not showing unique associations between performance and BOLD activity, namely the fusiform gyrus and the posterior cingulate cortex (Figure 4, orange regions). This is consistent with the view that differences in baseline CBF can affect the sign and the magnitude of the evoked BOLD signal, without affecting changes in the underlying neural activity (Cohen et al., 2002; Brown et al., 2003; Stefanovic et al., 2006). We extend prior findings by showing that only a portion of the CBF effects can introduce such a behaviourally irrelevant bias; other parts of the CBF variance might be related to behaviour-relevant signal, i.e. differences in CBF could be important in their own right. Unlike the normalisation approach described in Introduction to control for CBF differences, the current commonality framework allows partition of CBF effects into effects of interest and effects of no interest. We propose that modelling the effects of age on baseline CBF, and in general cerebrovascular and neurovascular health (Tsvetanov et al., 2021), has implications for the interpretation of fMRI studies of ageing, whereby it can improve brain-behaviour relationships and provide a viable mechanistic account of maintaining and improving cognitive function in old age.

### 4.3. Unique effects of performance, age and CBF on task-related activity

After accounting for age and performance, higher baseline cerebral blood flow remained significantly associated with the level of BOLD activity in cortical regions modulated by demanding problem-solving processes (Figure 3). Higher baseline CBF related to higher range of activation in task positive regions under more demanding processing, including the middle frontal gyrus, the putamen, and the cuneus. The effects were spatially adjacent or overlapping with behaviour-relevant region suggesting that higher baseline CBF may provide the conditions to upregulate activity in these regions, possibly through functional hyperaemia. Additionally, higher CBF provided higher range of deactivation in task negative regions, namely the angular gyrus and precentral gyrus. These effects were spatially adjacent or overlapping with regions showing inefficient deactivation with ageing and suggest that higher baseline CBF may facilitate suppression of activity in task-negative regions. This may reflect the effect of having an intact vasodilatory reserve (Sobczyk et al., 2014; Digernes et al., 2017). Our findings have direct implications for task-based BOLD imaging whereby higher baseline CBF levels contribute to stronger changes in BOLD signal amplitude in response to demanding cognitive conditions. The myogenic response and cardiac output are two major modulators of resting CBF (Hill et al., 2006; Meng et al., 2015), which require future consideration to establish the mechanism underlying our findings.

Ageing was associated with weaker activation of the multiple demand network and less efficient suppression of the default network. These effects were over and above performance and CBF, suggesting the involvement of additional factors leading to age-related difference in BOLD activity. Some factors include genetics (Shan et al., 2016), cardiovascular and neurovascular signals not captured by baseline CBF (Abdelkarim et al., 2019; Tsvetanov et al., 2021) or effects of functional connectivity captured by regional activity (Tsvetanov et al., 2018). Age differences in the shape of the haemodynamic response function (West et al., 2019) are less likely to introduce bias in the current study given its block-related fMRI design (Liu et al., 2001). The nature of these age effects should be elucidated through further investigation. The commonality analysis framework provides a useful tool for multivariate simultaneous modelling to disentangle the multifactorial nature of age-related BOLD differences.

After accounting for age, baseline CBF and other covariates of not interest, the level of activity in the multiple-demand regions remained positively associated with performance during the Cattell task in the scanner. Our findings are in line with previous studies during diverse demanding tasks, including manipulations of working memory, target detection, response inhibition (Fedorenko et al., 2013; Tschentscher et al., 2017; Assem et al., 2020a, 2020b). Given that both age and cerebrovascular reactivity could introduce a very strong effect on the activity-behaviour associations (even with narrow age range and healthy populations), our approach to control for these factors, in combination with the population-based, large-sample, provide the strongest evidence to date that individual differences variance in executive abilities is selectively and robustly associated with the level of activity in the multiple demand network.

Our study adds evidence to the nature of suppression of the default network during externally directed task (Buckner and DiNicola, 2019). The task-induced default network deactivations were consistent with previous findings in the Cattell task (Samu et al., 2017) and in general with the extent to which task conditions are cognitive demanding (Anticevic et al., 2012; Sripada et al., 2020). The effects in the default network were related to age or baseline CBF, but not uniquely related to performance, suggesting that the level of BOLD deactivations during Cattell task do not reflect individual variability in cognitive performance. The nature of default network suppression remains to be fully defined (Fox et al., 2018), but future findings about the default network cannot be interpreted independent of age and baseline CBF, at least when aiming to understand the relevance of DMN suppression in health and disease.

## 5. Issues and future directions

There are issues to the study. Our findings are based on a population-based cross-sectional cohort, which cannot directly speak to individual’s progression over time (i.e, the ageing process). We only assessed the brain activations/co-activations, but do not quantify brain connectivity (Tsvetanov et al., 2016, 2020a; Geerligs et al., 2017; Samu et al., 2017; Bethlehem et al., 2020), even though both may change with in cognitive ageing (Tsvetanov et al., 2018). The relationship between baseline CBF and functional connectivity decouples with ageing (Galiano et al., 2019), but the behavioural relevance of such decoupling remains unclear, albeit motivated by prior work controlling for vascular effects from fMRI BOLD data (Tsvetanov et al., 2016; Geerligs et al., 2017). Future work should also i) evaluate the effects of CBF under different cognitive states (Campbell et al., 2015; Geerligs and Tsvetanov, 2016), ii) consider nonlinearities between CBF and BOLD signal within individuals (Chen, 2019) and across the lifespan (Tsvetanov et al., 2016; Tibon et al., 2021), and iii) the relevance of baseline CBF to stimulus-evoked CBF (Jennings et al., 2005) and other measures of cerebrovascular reactivity in ageing (Tsvetanov et al., 2020b) and neurodegenerative diseases (Chen, 2019).

## 6. Conclusion

We introduce a novel approach to neuroimaging that can dissociate between shared and unique signals across multiple neuroimaging modalities. Using this method, we show the effects of age on cerebral blood flow, task-related BOLD responses and performance. The results demonstrate that cerebrovascular health (i.e., baseline cerebral blood flow) explains confounding but also performance-related BOLD responses in fluid ability across the lifespan. They highlight the importance of using resting CBF data to model, rather than simply normalise for, differences in vascular health in task-based fMRI BOLD data (cf. Tsvetanov et al., 2021). Unlike the normalisation approach, our approach allows simultaneous modelling of multiple measures with independent contributions to cerebrovascular health. Here, we provide empirical evidence in support of the mechanism underlying the link between baseline CBF and neurocognitive function across the lifespan. The insights from our results may facilitate the development of new strategies to maintain cognitive ability across the life span in health and disease.

## 7. Acknowledgements

This work is supported by the Guarantors of Brain (G101149), SCNU Study Abroad Program for Elite Postgraduate Students, the Medical Research Council (MC_UU_00005/12/SUAG004/051/RG91365; SUAG04/51 R101400) and the Cambridge NIHR Biomedical Research Centre (BRC-1215-20014). The views expressed are those of the authors and not necessarily those of the NIHR or the Department of Health and Social Care. For the purpose of open access, the author has applied a CC BY public copyright licence to any Author Accepted Manuscript version arising from this submission. The Cambridge Centre for Ageing and Neuroscience (Cam-CAN) research was supported by the Biotechnology and Biological Sciences Research Council (grant number BB/H008217/1). We thank the Cam-CAN respondents and their primary care teams in Cambridge for their participation in this study. Further information about the Cam-CAN corporate authorship membership can be found at https://www.cam-can.org/index.php?content=corpauth#13.

## References

Abdelkarim D, Zhao Y, Turner MP, Sivakolundu DK, Lu H, Rypma B (2019) A neural-vascular complex of age-related changes in the human brain: Anatomy, physiology, and implications for neurocognitive aging. Neurosci Biobehav Rev 107:927–944 Available at: http://www.ncbi.nlm.nih.gov/pubmed/31499083 [Accessed September 26, 2019].

Addicott MA, Yang LL, Peiffer AM, Burnett LR, Burdette JH, Chen MY, Hayasaka S, Kraft RA, Maldjian JA, Laurienti PJ (2009) The effect of daily caffeine use on cerebral blood flow: How much caffeine can we tolerate? Hum Brain Mapp 30:3102–3114.

Anticevic A, Cole MW, Murray JD, Corlett PR, Wang X-J, Krystal JH (2012) The role of default network deactivation in cognition and disease. Trends Cogn Sci 16:584–592 Available at: http://www.ncbi.nlm.nih.gov/pubmed/23142417 [Accessed August 2, 2013].

Assem M, Blank IA, Mineroff Z, Ademoğlu A, Fedorenko E (2020a) Activity in the fronto-parietal multiple-demand network is robustly associated with individual differences in working memory and fluid intelligence. Cortex 131:1–16.

Assem M, Glasser MF, Van Essen DC, Duncan J (2020b) A Domain-General Cognitive Core Defined in Multimodally Parcellated Human Cortex. Cereb Cortex 30:4361–4380 Available at: https://academic.oup.com/cercor/article/30/8/4361/5815289 [Accessed June 28, 2021].

Bangen KJ, Nation DA, Clark LR, Harmell AL, Wierenga CE, Dev SI, Delano-Wood L, Zlatar ZZ, Salmon DP, Liu TT, Bondi MW (2014) Interactive effects of vascular risk burden and advanced age on cerebral blood flow. Front Aging Neurosci 6:1–10.

Bethlehem RAI, Paquola C, Seidlitz J, Ronan L, Bernhardt B, Consortium C-C, Tsvetanov KA (2020) Dispersion of functional gradients across the adult lifespan. Neuroimage:117299 Available at: https://linkinghub.elsevier.com/retrieve/pii/S1053811920307850 [Accessed August 27, 2020].

Birn RM, Diamond JB, Smith M a, Bandettini P a (2006) Separating respiratory-variation-related fluctuations from neuronal-activity-related fluctuations in fMRI. Neuroimage 31:1536–1548 Available at: http://www.ncbi.nlm.nih.gov/pubmed/16632379 [Accessed February 28, 2013].

Brown GG, Eyler Zorrilla LT, Georgy B, Kindermann SS, Wong EC, Buxton RB (2003) BOLD and perfusion response to finger-thumb apposition after acetazolamide administration: differential relationship to global perfusion. J Cereb Blood Flow Metab 23:829–837 Available at: http://www.ncbi.nlm.nih.gov/pubmed/12843786 [Accessed October 1, 2019].

Buckner RL, Andrews-Hanna JR, Schacter DL (2008) The brain’s default network: anatomy, function, and relevance to disease. Ann N Y Acad Sci 1124:1–38 Available at: http://www.ncbi.nlm.nih.gov/pubmed/18400922 [Accessed May 21, 2013].

Buckner RL, DiNicola LM (2019) The brain’s default network: updated anatomy, physiology and evolving insights. Nat Rev Neurosci 20:593–608 Available at: www.nature.com/nrn [Accessed June 29, 2021].

Cabeza R, Albert M, Belleville S, Craik FIM, Duarte A, Grady CL, Lindenberger U, Nyberg L, Park DC, Reuter-Lorenz PA, Rugg MD, Steffener J, Rajah MN (2018) Maintenance, reserve and compensation: the cognitive neuroscience of healthy ageing. Nat Rev Neurosci 19:701–710 Available at: http://www.nature.com/articles/s41583-018-0068-2 [Accessed March 3, 2019].

Camilleri JA, Müller VI, Fox P, Laird AR, Hoffstaedter F, Kalenscher T, Eickhoff SB (2018) Definition and characterization of an extended multiple-demand network. Neuroimage 165:138–147 Available at: https://pubmed.ncbi.nlm.nih.gov/29030105/ [Accessed December 15, 2020].

Campbell KL et al. (2015) Idiosyncratic responding during movie-watching predicted by age differences in attentional control. Neurobiol Aging 36:3045–3055.

Cattell RB (1971) Abilities: Their structure growth and action. Boston, MA: Houghton Mifflin.

Chen JJ (2019) Functional MRI of brain physiology in aging and neurodegenerative diseases. Neuroimage 187:209–225.

Chen JJ, Rosas HD, Salat DH (2011) Age-associated reductions in cerebral blood flow are independent from regional atrophy. Neuroimage 55:468–478 Available at: http://www.sciencedirect.com/science/article/pii/S1053811910016162 [Accessed March 21, 2014].

Cohen ER, Ugurbil K, Kim S-G (2002) Effect of Basal Conditions on the Magnitude and Dynamics of the Blood Oxygenation Level-Dependent fMRI Response. J Cereb Blood Flow Metab 22:1042–1053 Available at: http://www.ncbi.nlm.nih.gov/pubmed/12218410 [Accessed September 29, 2019].

Crittenden BM, Mitchell DJ, Duncan J (2016) Task encoding across the multiple demand cortex is consistent with a frontoparietal and cingulo-opercular dual networks distinction. J Neurosci 36:6147–6155 Available at: https://www.jneurosci.org/content/36/23/6147 [Accessed December 15, 2020].

Digernes I, Bjørnerud A, Vatnehol SAS, Løvland G, Courivaud F, Vik-Mo E, Meling TR, Emblem KE (2017) A theoretical framework for determining cerebral vascular function and heterogeneity from dynamic susceptibility contrast MRI: J Cereb Blood Flow Metab 37:2237–2248 Available at: https://journals.sagepub.com/doi/full/10.1177/0271678X17694187 [Accessed August 13, 2021].

Domino EF, Ni L, Xu Y, Koeppe RA, Guthrie S, Zubieta JK (2004) Regional cerebral blood flow and plasma nicotine after smoking tobacco cigarettes. Prog Neuro-Psychopharmacology Biol Psychiatry 28:319–327.

Duncan J (2010) The multiple-demand (MD) system of the primate brain: mental programs for intelligent behaviour. Trends Cogn Sci 14:172–179 Available at: http://www.ncbi.nlm.nih.gov/pubmed/20171926 [Accessed October 17, 2013].

Duncan J (2013) The structure of cognition: attentional episodes in mind and brain. Neuron 80:35–50.

Fedorenko E, Duncan J, Kanwisher N (2013) Broad domain generality in focal regions of frontal and parietal cortex. Proc Natl Acad Sci U S A 110:16616–16621 Available at: https://pubmed.ncbi.nlm.nih.gov/24062451/ [Accessed May 19, 2021].

Folstein MF, Folstein SE, McHugh PR (1975) “Mini-mental state.” J Psychiatr Res 12:189–198 Available at: http://www.sciencedirect.com/science/article/pii/0022395675900266 [Accessed February 21, 2014].

Fox K, Foster B, Kucyi A, Daitch A, Parvizi J (2018) Intracranial Electrophysiology of the Human Default Network. Trends Cogn Sci 22:307–324 Available at: https://pubmed.ncbi.nlm.nih.gov/29525387/ [Accessed August 20, 2021].

Friston KJ, Ashburner J, Kiebel S, Nichols T, Penny WD (2007) Statistical parametric mapping : the analysis of funtional brain images. Elsevier Academic Press.

Gaballa MA, Jacob CT, Raya TE, Liu J, Simon B, Goldman S (1998) Large Artery Remodeling During Aging. Hypertension 32:437–443 Available at: https://www.ahajournals.org/doi/abs/10.1161/01.HYP.32.3.437 [Accessed August 20, 2021].

Galiano A, Mengual E, García de Eulate R, Galdeano I, Vidorreta M, Recio M, Riverol M, Zubieta JL, Fernández-Seara MA (2019) Coupling of cerebral blood flow and functional connectivity is decreased in healthy aging. Brain Imaging Behav:1–15 Available at: http://link.springer.com/10.1007/s11682-019-00157-w [Accessed September 26, 2019].

Geerligs L, Tsvetanov KA (2016) The use of resting state data in an integrative approach to studying neurocognitive ageing – Commentary on Campbell and Schacter (2016). Lang Cogn Neurosci 32:684–691.

Geerligs L, Tsvetanov KA, Cam-Can, Henson RN (2017) Challenges in measuring individual differences in functional connectivity using fMRI: The case of healthy aging. Hum Brain Mapp.

Grade M, Hernandez Tamames JA, Pizzini FB, Achten E, Golay X, Smits M (2015) A neuroradiologist’s guide to arterial spin labeling MRI in clinical practice. Neuroradiology 57:1181–1202.

Hays CC, Zlatar ZZ, Campbell L, Meloy MJ, Wierenga CE (2017) Temporal gradient during famous face naming is associated with lower cerebral blood flow and gray matter volume in aging. Neuropsychologia 107:76–83 Available at: https://pubmed.ncbi.nlm.nih.gov/29133109/ [Accessed July 11, 2020].

Henriksen OM, Vestergaard MB, Lindberg U, Aachmann-Andersen NJ, Lisbjerg K, Christensen SJ, Rasmussen P, Olsen N V., Forman JL, Larsson HBW, Law I (2018) Interindividual and regional relationship between cerebral blood flow and glucose metabolism in the resting brain. https://doi.org/101152/japplphysiol002762018 125:1080–1089 Available at: https://journals.physiology.org/doi/abs/10.1152/japplphysiol.00276.2018 [Accessed September 22, 2021].

Hill MA, Davis MJ, Meininger GA, Potocnik SJ, Murphy T V. (2006) Arteriolar myogenic signalling mechanisms: Implications for local vascular function. Clin Hemorheol Microcirc 34:67–79.

Horn JL, Cattell RB (1967) Age differences in fluid and crystallized intelligence. Acta Psychol (Amst) 26:107–129.

Hutchison JL, Lu H, Rypma B (2013) Neural Mechanisms of Age-Related Slowing: The ΔCBF/ΔCMRO2 Ratio Mediates Age-Differences in BOLD Signal and Human Performance. Cereb cortex 23:2337–2346 Available at: http://www.ncbi.nlm.nih.gov/pubmed/22879349 [Accessed November 27, 2012].

Iadecola C (2004) Neurovascular regulation in the normal brain and in Alzheimer’s disease. Nat Rev Neurosci 5:347–360 Available at: http://www.nature.com/doifinder/10.1038/nrn1387 [Accessed August 15, 2017].

Jennings JR, Muldoon MF, Ryan C, Price JC, Greer P, Sutton-Tyrrell K, Veen FM van der, Meltzer CC (2005) Reduced cerebral blood flow response and compensation among patients with untreated hypertension. Neurology 64:1358–1365 Available at: https://n.neurology.org/content/64/8/1358 [Accessed August 21, 2021].

Kaufman AS, Horn JL (1996) Age changes on tests of fluid and crystallized ability for women and men on the Kaufman Adolescent and Adult Intelligence Test (KAIT) at ages 17-94 years. Arch Clin Neuropsychol 11:97–121.

Kennedy KM, Raz N (2015) Normal Aging of the Brain. In: Brain Mapping, pp 603–617. Elsevier. Available at: https://linkinghub.elsevier.com/retrieve/pii/B9780123970251000683 [Accessed February 5, 2019].

Kievit RA et al. (2014) Distinct aspects of frontal lobe structure mediate age-related differences in fluid intelligence and multitasking. Nat Commun 5:5658 Available at: http://www.nature.com/doifinder/10.1038/ncomms6658 [Accessed September 13, 2017].

Kisler K, Nelson AR, Montagne A, Zlokovic B V. (2017) Cerebral blood flow regulation and neurovascular dysfunction in Alzheimer disease. Nat Rev Neurosci 18:419–434 Available at: http://www.ncbi.nlm.nih.gov/pubmed/28515434 [Accessed August 18, 2017].

Kraha A, Turner H, Nimon K, Zientek LR, Henson RK (2012) Tools to Support Interpreting Multiple Regression in the Face of Multicollinearity. Front Psychol 3:44 Available at: http://journal.frontiersin.org/article/10.3389/fpsyg.2012.00044/abstract [Accessed August 22, 2017].

Leeuwis AE, Smith LA, Melbourne A, Hughes AD, Richards M, Prins ND, Sokolska M, Atkinson D, Tillin T, Jäger HR, Chaturvedi N, Flier WM van der, Barkhof F (2018) Cerebral Blood Flow and Cognitive Functioning in a Community-Based, Multi-Ethnic Cohort: The SABRE Study. Front Aging Neurosci 10:279 Available at: https://www.frontiersin.org/article/10.3389/fnagi.2018.00279/full [Accessed December 17, 2020].

Lemkuil BP, Drummond JC, Patel PM (2013) Central Nervous System Physiology: Cerebrovascular. Pharmacol Physiol Anesth Found Clin Appl:123–136.

Li Y, Shen Q, Huang S, Li W, Muir E, Long J, TQ D (2015) Cerebral angiography, blood flow and vascular reactivity in progressive hypertension. Neuroimage 111:329–337 Available at: https://pubmed.ncbi.nlm.nih.gov/25731987/ [Accessed August 20, 2021].

Liu P, Hebrank AC, Rodrigue KM, Kennedy KM, Section J, Park DC, Lu H (2013) Age-related differences in memory-encoding fMRI responses after accounting for decline in vascular reactivity. Neuroimage 78:415–425 Available at: http://www.ncbi.nlm.nih.gov/pubmed/23624491 [Accessed October 24, 2013].

Liu TT, Frank LR, Wong EC, Buxton RB (2001) Detection Power, Estimation Efficiency, and Predictability in Event-Related fMRI. Neuroimage 13:759–773.

Meng L, Hou W, Chui J, Han R, Gelb A (2015) Cardiac Output and Cerebral Blood Flow: The Integrated Regulation of Brain Perfusion in Adult Humans. Anesthesiology 123:1198–1208 Available at: https://pubmed.ncbi.nlm.nih.gov/26402848/ [Accessed August 13, 2021].

Merola A, Germuska MA, Warnert EA, Richmond L, Helme D, Khot S, Murphy K, Rogers PJ, Hall JE, Wise RG (2017) Mapping the pharmacological modulation of brain oxygen metabolism: The effects of caffeine on absolute CMRO2 measured using dual calibrated fMRI. Neuroimage 155:331–343 Available at: https://www.sciencedirect.com/science/article/pii/S1053811917302367?via%3Dihub [Accessed October 7, 2019].

Mishra A, Hall CN, Howarth C, Freeman RD (2021) Key relationships between non-invasive functional neuroimaging and the underlying neuronal activity. Philos Trans R Soc B Biol Sci 376:20190622 Available at: https://royalsocietypublishing.org/doi/10.1098/rstb.2019.0622 [Accessed December 17, 2020].

Mutsaerts HJ, Petr J, Václavů L, van Dalen JW, Robertson AD, Caan MW, Masellis M, Nederveen AJ, Richard E, MacIntosh BJ (2017) The spatial coefficient of variation in arterial spin labeling cerebral blood flow images. J Cereb Blood Flow Metab 37:3184–3192 Available at: http://www.ncbi.nlm.nih.gov/pubmed/28058975 [Accessed June 25, 2019].

Mutsaerts HJMM et al. (2018) Comparison of arterial spin labeling registration strategies in the multi-center GENetic frontotemporal dementia initiative (GENFI). J Magn Reson Imaging 47:131–140 Available at: http://www.ncbi.nlm.nih.gov/pubmed/28480617 [Accessed September 16, 2019].

Nimon K, Lewis M, Kane R, Haynes RM (2008) An R package to compute commonality coefficients in the multiple regression case: An introduction to the package and a practical example. Behav Res Methods 40:457–466.

Ohanian J, Liao A, Forman SP, Ohanian V (2014) Age-related remodeling of small arteries is accompanied by increased sphingomyelinase activity and accumulation of long-chain ceramides. Physiol Rep 2 Available at: /pmc/articles/PMC4098743/ [Accessed August 20, 2021].

Patricia C, Henk-Jan M, Eidrees G, Marion S, Marjan A, Egill R, Francesca Benedetta P, Jorge J, Mervi K, Ritva V, António B-L, Roland W, Elna-Marie L, Eric A (2014) Review of confounding effects on perfusion measurements. Front Hum Neurosci 8 Available at: http://www.frontiersin.org/Community/AbstractDetails.aspx?ABS_DOI=10.3389/conf.fnhum.2014.214.00073 [Accessed October 4, 2019].

Piguet O, Hornberger M, Shelley BP, Kipps CM, Hodges JR (2009) Sensitivity of current criteria for the diagnosis of behavioral variant frontotemporal dementia. Neurology 72:732–737.

Raichle ME (2015) The Brain’s Default Mode Network. Annu Rev Neurosci:413–427.

Rorden C, Brett M (2000) Stereotaxic display of brain lesions. Behav Neurol 12:191–200 Available at: https://pubmed.ncbi.nlm.nih.gov/11568431/ [Accessed November 8, 2021].

Salthouse T (2012) Consequences of age-related cognitive declines. Annu Rev Psychol 63:201–226 Available at: http://www.ncbi.nlm.nih.gov/pubmed/21740223 [Accessed November 27, 2012].

Salthouse TA, Atkinson TM, Berish DE (2003) Executive functioning as a potential mediator of age-related cognitive decline in normal adults. J Exp Psychol Gen 132:566–594.

Samu D et al. (2017) Preserved cognitive functions with age are determined by domain-dependent shifts in network responsivity. Nat Commun 8:ncomms14743.

Shafto MA et al. (2014) The Cambridge Centre for Ageing and Neuroscience (Cam-CAN) study protocol: A cross-sectional, lifespan, multidisciplinary examination of healthy cognitive ageing. BMC Neurol 14.

Shan ZY, Vinkhuyzen AAE, Thompson PM, McMahon KL, Blokland GAM, de Zubicaray GI, Calhoun V, Martin NG, Visscher PM, Wright MJ, Reutens DC (2016) Genes influence the amplitude and timing of brain hemodynamic responses. Neuroimage 124:663–671.

Shmuel A, Yacoub E, Pfeuffer J, Van de Moortele PF, Adriany G, Hu X, Ugurbil K (2002) Sustained negative BOLD, blood flow and oxygen consumption response and its coupling to the positive response in the human brain. Neuron 36:1195–1210.

Skinner HA (1982) The drug abuse screening test. Addict Behav 7:363–371.

Smith SM, Nichols TE (2009) Threshold-free cluster enhancement: Addressing problems of smoothing, threshold dependence and localisation in cluster inference. Neuroimage 44:83–98 Available at: https://pubmed.ncbi.nlm.nih.gov/18501637/ [Accessed June 21, 2021].

Snellen H (1862) Probebuchstaben zur bestimmung der sehscharfe. Utrecht: Van de Weijer.

Sobczyk O, Battisti-Charbonney a, Fierstra J, Mandell DM, Poublanc J, Crawley a P, Mikulis DJ, Duffin J, Fisher J a (2014) A conceptual model for CO2-induced redistribution of cerebral blood flow with experimental confirmation using BOLD MRI. Neuroimage 92C:56–68 Available at: http://www.ncbi.nlm.nih.gov/pubmed/24508647 [Accessed March 22, 2014].

Sripada C, Angstadt M, Rutherford S, Taxali A, Shedden K (2020) Toward a “treadmill test” for cognition: Improved prediction of general cognitive ability from the task activated brain. Hum Brain Mapp 41:3186–3197 Available at: https://onlinelibrary.wiley.com/doi/full/10.1002/hbm.25007 [Accessed June 28, 2021].

Stefanovic B, Warnking JM, Rylander KM, Pike GB (2006) The effect of global cerebral vasodilation on focal activation hemodynamics. Neuroimage 30:726–734.

Sweeney MD, Kisler K, Montagne A, Toga AW, Zlokovic B V. (2018) The role of brain vasculature in neurodegenerative disorders. Nat Neurosci 21:1318–1331.

Sweeney MD, Zhao Z, Montagne A, Nelson AR, Zlokovic B V. (2019) Blood-brain barrier: From physiology to disease and back. Physiol Rev 99:21–78.

Taylor JR, Williams N, Cusack R, Auer T, Shafto MA, Dixon M, Tyler LK, Cam-Can, Henson RN (2015) The Cambridge Centre for Ageing and Neuroscience (Cam-CAN) data repository: Structural and functional MRI, MEG, and cognitive data from a cross-sectional adult lifespan sample. Neuroimage Available at: http://www.sciencedirect.com/science/article/pii/S1053811915008150 [Accessed September 21, 2015].

Tibon R, Tsvetanov KA, Price D, Nesbitt D, Can C, Henson R (2021) Transient neural network dynamics in cognitive ageing. Neurobiol Aging 105:217–228.

Tschentscher N, Mitchell D, Duncan J (2017) Fluid intelligence predicts novel rule implementation in a distributed frontoparietal control network. J Neurosci 37:4841–4847.

Tsvetanov KA et al. (2020a) Brain functional network integrity sustains cognitive function despite atrophy in presymptomatic genetic frontotemporal dementia. Alzheimer’s Dement:alz.12209 Available at: https://onlinelibrary.wiley.com/doi/10.1002/alz.12209 [Accessed December 9, 2020].

Tsvetanov KA, Henson RNA, Jones PS, Mutsaerts H, Fuhrmann D, Tyler LK, Rowe JB (2020b) The effects of age on resting-state BOLD signal variability is explained by cardiovascular and cerebrovascular factors. In: Psychophysiology. Blackwell Publishing Inc. Available at: https://onlinelibrary.wiley.com/doi/full/10.1111/psyp.13714 [Accessed December 9, 2020].

Tsvetanov KA, Henson RNA, Rowe JB (2021) Separating vascular and neuronal effects of age on fMRI BOLD signals. Philos Trans R Soc London Ser B, Biol Sci 376:20190631.

Tsvetanov KA, Henson RNA, Tyler LK, Davis SW, Shafto MA, Taylor JR, Williams N, Rowe JB (2015) The effect of ageing on fMRI: Correction for the confounding effects of vascular reactivity evaluated by joint fMRI and MEG in 335 adults. Hum Brain Mapp 36:2248–2269 Available at: http://www.ncbi.nlm.nih.gov/pubmed/25727740 [Accessed February 27, 2015].

Tsvetanov KA, Henson RNA, Tyler LK, Razi A, Geerligs L, Ham TE, Rowe JB (2016) Extrinsic and intrinsic brain network connectivity maintains cognition across the lifespan despite accelerated decay of regional brain activation. J Neurosci 36:3115–3126.

Tsvetanov KA, Ye Z, Hughes L, Samu D, Treder MS, Wolpe N, Tyler LK, Rowe JB, for Cambridge Centre for Ageing and Neuroscience (2018) Activity and connectivity differences underlying inhibitory control across the adult lifespan. J Neurosci 38:7887–7900 Available at: http://www.ncbi.nlm.nih.gov/pubmed/30049889 [Accessed August 1, 2018].

United Nations D of E and SAPD (2020) World Population Ageing 2019.

West KL, Zuppichini MD, Turner MP, Sivakolundu DK, Zhao Y, Abdelkarim D, Spence JS, Rypma B (2019) BOLD hemodynamic response function changes significantly with healthy aging. Neuroimage 188:198–207.

Willie CK, Tzeng Y-C, Fisher JA, Ainslie PN (2014) Integrative regulation of human brain blood flow. J Physiol 592:841–859 Available at: http://www.ncbi.nlm.nih.gov/pubmed/24396059 [Accessed October 1, 2019].

Woolgar A, Bor D, Duncan J (2013) Global increase in task-related fronto-parietal activity after focal frontal lobe lesion. J Cogn Neurosci 25:1542–1552.

Woolgar A, Duncan J, Manes F, Fedorenko E (2018) Fluid intelligence is supported by the multiple-demand system not the language system. Nat Hum Behav 2:200–204 Available at: https://pubmed.ncbi.nlm.nih.gov/31620646/ [Accessed December 15, 2020].

Yarchoan M, Xie SX, Kling MA, Toledo JB, Wolk DA, Lee EB, Van Deerlin V, Lee VMY, Trojanowski JQ, Arnold SE (2012) Cerebrovascular atherosclerosis correlates with Alzheimer pathology in neurodegenerative dementias. Brain 135:3749–3756.

Zhang N, Gordon ML, Ma Y, Chi B, Gomar JJ, Peng S, Kingsley PB, Eidelberg D, Goldberg TE (2018) The Age-Related Perfusion Pattern Measured With Arterial Spin Labeling MRI in Healthy Subjects. Front Aging Neurosci 10:214 Available at: http://www.ncbi.nlm.nih.gov/pubmed/30065646 [Accessed July 9, 2019].

Zientek LR, Thompson B (2006) Commonality analysis: Partitioning variance to facilitate better understanding of data. J Early Interv 28:299–307 Available at: https://journals.sagepub.com/doi/abs/10.1177/105381510602800405?journalCode=jeib [Accessed June 21, 2021].

Zlokovic B V (2011) Neurovascular pathways to neurodegeneration in Alzheimer’s disease and other disorders. Nat Rev Neurosci 12:723–738 Available at: http://www.ncbi.nlm.nih.gov/pubmed/22048062 [Accessed August 19, 2017].

